# Copper Import via CTR1 Supports the β3-Adrenergic Thermogenic Program

**DOI:** 10.64898/2026.03.24.713962

**Authors:** Tae-Il Jeon, Young-Seung Lee, Tamara Korolnek, Juyoung Kim, Pratibha Poudel, Parama Bhattacharjee, Xuewei Zhao, Evan Ying, Naili Liu, Tong Xiao, Christopher J. Chang, Oksana Gavrilova, Byung-Eun Kim

## Abstract

Adaptive thermogenesis requires coordinated activation of mitochondrial oxidation and metabolic remodeling, yet the signals driving this coordination are incompletely understood. Here, we show that cold exposure and β3-adrenergic receptor (β3-AR) stimulation upregulate the high-affinity copper (Cu) importer CTR1 and promote Cu accumulation in thermogenic adipose tissues. Adipocyte-specific *Ctr1* knockout (ACKO) mice exhibit markedly reduced energy expenditure and develop severe hypothermia during acute cold challenge. Proteomic analysis of brown adipose tissue (BAT) from ACKO mice reveals coordinated suppression of oxidative phosphorylation and thermogenic metabolic programs, accompanied by attenuation of lipolytic pathways. Cu deficiency also impairs cold- and β3-AR-induced lipolytic activation, including reduced HSL phosphorylation and lipid clearance in both BAT and inguinal white adipose tissue (iWAT). Although BAT-specific *Ctr1* deletion (BCKO) leaves acute β3-adrenergic responses largely intact, these mice still exhibit cold intolerance, indicating that BAT Cu homeostasis is indispensable for sustaining thermogenic capacity during cold challenge. Treatment with the Cu ionophore elesclomol partially restores mitochondrial oxidative capacity and improves cold tolerance in ACKO mice. Together, these findings identify CTR1-dependent Cu import as a dynamically regulated component of the β3-adrenergic thermogenic program and establish intracellular Cu availability as a key determinant of thermogenic capacity during adaptive thermogenesis.

## INTRODUCTION

Copper (Cu) is an essential trace element required for normal growth and development^1^. Mutations disrupting Cu homeostasis cause severe inherited disorders, including Wilson’s disease and Menkes disease²; dysregulated Cu metabolism has also been linked to neurodegeneration and cardiomyopathy^2^. Cellular Cu uptake occurs primarily through the high-affinity importer CTR1, followed by intracellular capture by the metalloadaptor RAD23B^3^, which serves as a hub for distribution to specific intracellular targets via dedicated metallochaperones. The metallochaperone CCS delivers Cu to Cu/Zn superoxide dismutase (SOD1), whereas ATOX1 transfers Cu to the P-type ATPases ATP7A and ATP7B, which regulate Cu export and secretory pathway trafficking^4, 5^. The physiological importance of this network is exemplified by Menkes disease, caused by ATP7A mutations that result in systemic Cu deficiency with connective tissue abnormalities, severe neurodegeneration, and impaired thermogenesis^6, 7^. Beyond genetic causes, Cu deficiency can also arise from malabsorption, inadequate dietary intake, hyperzincemia, or bariatric surgery^8^. Although the molecular machinery governing cellular Cu transport has been extensively characterized, it remains unclear whether Cu import is dynamically regulated in specific tissues to support specialized metabolic programs or to meet increased metabolic demands, such as those occurring in adipose tissue during adaptive thermogenesis^9^.

Adaptive thermogenesis is a critical metabolic response to cold exposure. In thermogenic adipocytes, norepinephrine released from sympathetic nerves activates β3-adrenergic receptors (β3-AR), stimulating adenylyl cyclase, increasing cyclic AMP (cAMP), and activating protein kinase A (PKA)^10, 11^. This signaling cascade induces a thermogenic gene program via PGC-1α and cAMP-response element-binding protein (CREB) and promotes lipolysis through phosphorylation of hormone-sensitive lipase (HSL), releasing free fatty acids that fuel mitochondrial oxidation and activate uncoupling protein 1 (UCP1)^12, 13^. UCP1 dissipates the proton gradient generated by the electron transport chain (ETC), converting chemical energy into heat. Brown adipose tissue (BAT), enriched in mitochondria and constitutively expressing UCP1, responds rapidly to cold, whereas prolonged adrenergic stimulation induces “browning” of white adipose tissue (WAT), generating beige adipocytes with inducible thermogenic capacity^14^.

Mitochondrial respiratory capacity depends on adequate intracellular Cu availability, primarily for cytochrome c oxidase (complex IV), the terminal enzyme of the ETC that requires Cu as an essential catalytic cofactor. During cold exposure, thermogenic adipocytes markedly increase mitochondrial respiration, raising the possibility that Cu supply must be coordinated with thermogenic remodeling to sustain maximal respiratory flux. Supporting this possibility, hypothermia has been reported in some patients with Menkes disease^6, 15^. Although ATP7A deficiency impairs Cu delivery to dopamine β-hydroxylase in sympathetic neurons, potentially disrupting norepinephrine synthesis ^16, 17^, adipocyte-intrinsic Cu deficiency in Menkes patients may represent an additional, and potentially predominant, contributor to impaired thermogenesis. Consistent with a role for Cu in thermogenic adipocytes, Li and colleagues showed that Cu content increases in BAT during cold exposure^9^ and the Chang laboratory observed that Cu regulates lipolysis in WAT through metalloallostery^18^. However, the mechanisms underlying this increase and whether the Cu homeostasis machinery actively contributes to thermogenic activation remain unclear. In particular, it remains unknown whether Cu import is dynamically regulated during cold stress, whether intracellular Cu becomes limiting during adrenergic stimulation, or whether adipocyte-intrinsic Cu deficiency directly impairs mitochondrial function and lipolytic signaling downstream of β3-AR activation in vivo.

Here, we investigate the role of CTR1-mediated Cu import in adaptive thermogenesis using dietary Cu restriction, pan-adipocyte- and BAT-specific *Ctr1* knockout models, and pharmacologic Cu delivery. To determine whether impaired thermogenesis reflects reduced intracellular Cu availability, we employ the Cu ionophore elesclomol, which facilitates Cu transport independently of CTR1^19^. Together, these approaches reveal a previously unrecognized requirement for Cu homeostasis in the coordination of lipolytic activation and thermogenic remodeling of adipose tissue.

## 2. RESULTS

### 2.1. Systemic Cu deficiency impairs thermogenic adaptation to cold

To assess the impact of systemic Cu deficiency on the thermogenic response in wild-type (WT) mice, we evaluated cold tolerance using a dietary Cu-deficient model. WT mice (C57BL/6J) were maintained on a Cu-deficient (Cu-D) diet beginning at postnatal day 1 together with their mothers and continued on the same diet for 4 weeks after weaning (total duration of ∼7 weeks). Control Cu-adequate mice (Cu-A) received the same base chow but were supplemented with 20 mg L⁻¹ CuSO₄ in their drinking water.

Body weight was comparable between groups (Figure S1a) following the dietary intervention, indicating that the level and duration of dietary Cu restriction used in this study did not overtly impair growth. Cu-D mice exhibited characteristic features of Cu deficiency, including a pale coat (Figure S1b) and increased expression of the Cu chaperone for superoxide dismutase (CCS) in BAT (Figure 1a) and liver (Figure S1c). CCS expression level is inversely correlated with cytosolic Cu levels^20-22^, consistent with reduced tissue and cellular Cu availability in Cu-D mice.

**Figure 1.**
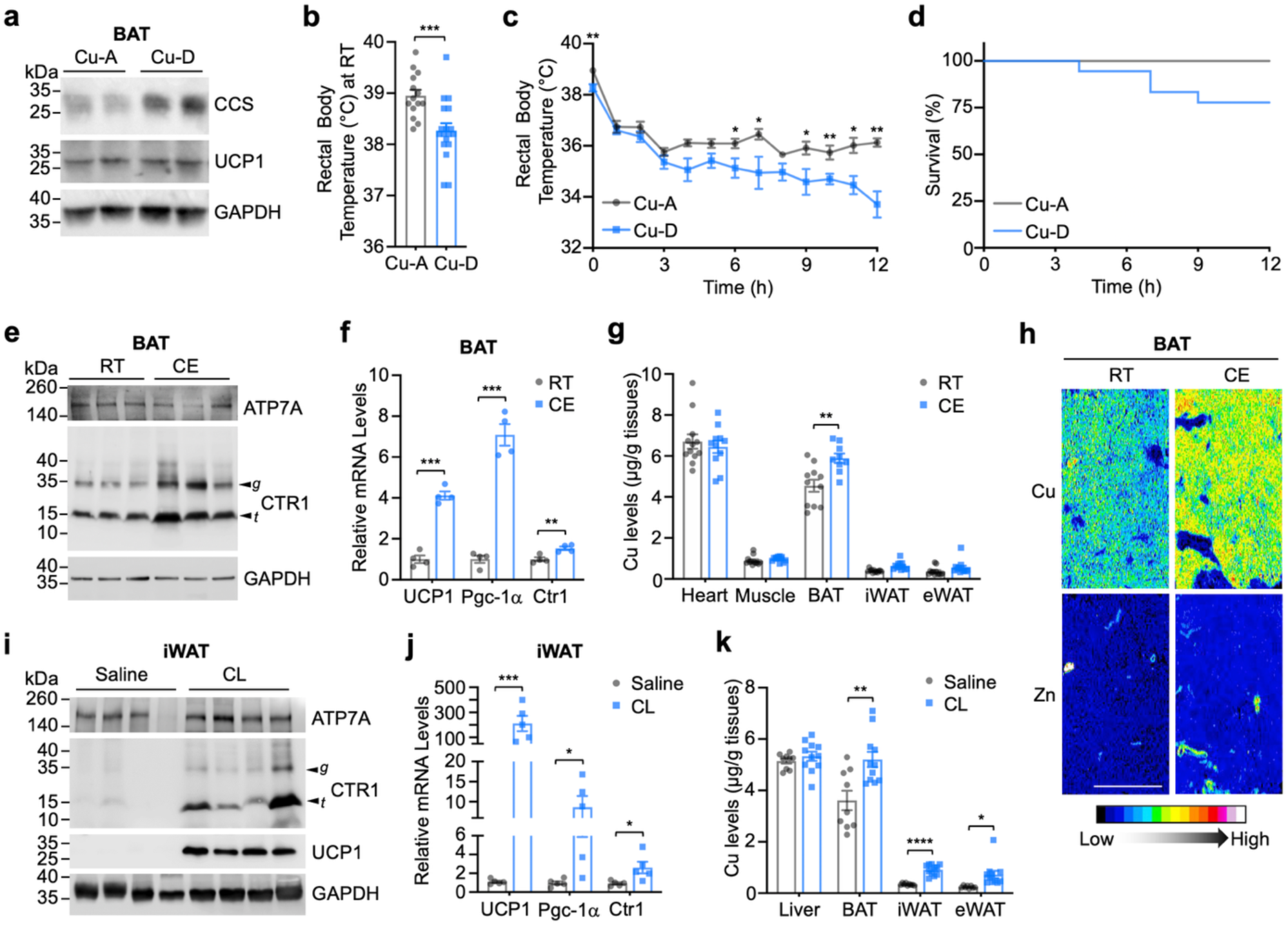
Cu availability is required for adaptive thermogenesis and is dynamically increased in thermogenic adipose tissue. (a) Immunoblot analysis of CCS and UCP1 protein levels in brown adipose tissue (BAT) from Cu-adequate (Cu-A) and Cu-deficient (Cu-D) mice. GAPDH served as a loading control. (b, c) Rectal body temperature of Cu-A (n = 14) and Cu-D (n = 18) mice maintained at (b) room temperature (RT) or exposed to (c) acute cold (CE, 4°C) for 12 h. (d) Kaplan–Meier survival analysis during 12 h of cold exposure (CE) for Cu-A (n = 14) and Cu-D (n = 18) mice (log-rank test, P = 0.647). (e) Immunoblot analysis of ATP7A and CTR1 in BAT from WT mice housed at RT or exposed to cold (CE, 4°C) for 12 h. Arrowheads indicate glycosylated full-length (g) and truncated (t) CTR1 species. (f) Relative mRNA expression of *Ucp1*, *Ppargc1a*, and *Slc31a1* (*Ctr1*) in BAT under RT and CE conditions (n = 4). (g) ICP–MS quantification of Cu levels in tissues from WT mice housed at RT or CE (n = 11 per condition). (h) Representative Laser Ablation (LA)–ICP–MS maps of Cu and Zn distribution in BAT from male WT mice housed at RT or CE (10 h). Scale bar, 500 μm. (i) Immunoblot analysis of ATP7A, CTR1, and UCP1 in inguinal white adipose tissue (iWAT) from WT mice treated with saline or CL316,243 (CL; 1 mg kg⁻¹ day⁻¹, i.p., 10 days). (j) Relative mRNA expression of *Ucp1*, *Ppargc1a*, and *Slc31a1* (*Ctr1*) in iWAT following saline or CL treatment (n = 5). (k) ICP–MS analysis of Cu levels in tissues from saline-treated (n = 9) or CL-treated (n = 11) mice. Unless otherwise indicated, mice were analyzed as mixed sex with balanced male and female representation across groups. Data are presented as mean ± SEM. Statistical significance was determined by two-tailed Student’s t-test. *P < 0.05, **P < 0.01, ***P < 0.001.

At room temperature (RT), Cu-D mice exhibited a modest but significant reduction in basal rectal temperature compared with Cu-A controls (Figure 1b). During acute cold exposure (4°C), both groups initially showed a drop in core temperature; however, Cu-A mice stabilized above ∼35°C, whereas Cu-D mice exhibited a progressive decline throughout the 12-h challenge (Figure 1c). Approximately 25% of Cu-D mice developed severe hypothermia during the 12-h cold challenge (Figure 1d). Although overall survival was not significantly different under this mildly Cu-deficient condition, the progressive temperature decline and the proportion of Cu-D mice reaching hypothermic thresholds support a role for Cu in thermoregulatory capacity during cold stress.

We next asked whether Cu handling is dynamically regulated during adaptive thermogenesis in WT mice housed on regular chow. After 12 h of cold exposure, protein levels of the Cu exporter ATP7A in BAT were unchanged, whereas the high-affinity Cu importer CTR1 (encoded by *Slc31a1*) was robustly increased (Figure 1e). Both glycosylated full-length and truncated CTR1 species were elevated, consistent with enhanced Cu import capacity^23^. Cold exposure markedly induced *Ucp1* and *Ppargc1a* (PGC-1α) expression, confirming activation of the thermogenic program, and *Slc31a1* (CTR1) mRNA was modestly but significantly increased (Figure 1f).

Inductively coupled plasma–mass spectrometry (ICP–MS) revealed a selective increase in total Cu content in BAT after cold exposure, with minimal changes in other tissues, including iWAT and epididymal white adipose tissue (eWAT) (Figure 1g), suggesting that thermogenic activation selectively increases Cu uptake in BAT. When analyzed separately by sex (Figure S1d), the increase in BAT Cu did not reach statistical significance in males; however, both female and male mice exhibited a consistent upward trend following cold exposure. Laser ablation (LA)–ICP–MS further demonstrated increased Cu signal throughout BAT, indicating a tissue-wide rise in Cu during cold challenge, but not zinc (Zn) (Figure 1h). Iron (Fe) levels also changed across tissues under cold conditions (Figure S1e), consistent with the known importance of Fe for mitochondrial biogenesis during adaptive thermogenesis^24-26^. These data suggest that adaptive thermogenesis is accompanied by increased Cu accumulation in BAT, likely driven by enhanced CTR1-dependent Cu import rather than decreased Cu export.

Cold exposure elicits multilayered metabolic remodeling across tissues through coordinated adrenergic and non-adrenergic pathways ^27-30^. Although systemic cold adaptation involves endocrine, immune, and metabolic rewiring, β3-adrenergic signaling in adipocytes represents a principal driver of thermogenic activation in rodents^27, 28^. To determine whether sympathetic activation is sufficient to induce Cu remodeling in adipose tissue, we selectively stimulated the β3-AR using CL316,243 (CL) for 10 days to induce browning of inguinal white adipose tissue (iWAT). CL treatment robustly increased UCP1 protein and mRNA levels in iWAT (Figure 1i,j), confirming activation of the thermogenic program. CTR1 protein abundance was markedly elevated, including both glycosylated and truncated species, whereas *Ctr1* mRNA increased only modestly (Figure 1i,j), suggesting that adrenergic signaling regulates CTR1 largely at the post-transcriptional level.

ICP–MS analysis revealed significant increases in Cu content in iWAT and BAT following CL treatment, with minimal changes in other tissues (Figure 1k). Fe levels were modestly altered in select depots (Figure S1f). Collectively, these results identify CTR1 induction and elevated Cu accumulation as integral components of thermogenic adipose remodeling induced by either cold exposure or β3-AR stimulation, suggesting increased Cu demand in thermogenic adipose tissues during thermogenic activation.

### 2.2. Adipocyte CTR1 is required for thermogenic adaptation

To directly test the physiological role of adipocyte CTR1, we generated adipose-specific *Slc31a1* (*Ctr1*) knockout mice (ACKO) by crossing *Ctr1*-floxed mice with *Adipoq*-Cre transgenic mice (Figure S2a,b). CTR1 protein was efficiently depleted in adipose depots (BAT and iWAT) but not in the liver (Figure 2a). Loss of CTR1 resulted in marked Cu deficiency in adipose tissue, as evidenced by increased CCS protein levels and a significant reduction in total Cu content in BAT and iWAT (Figure 2a,b). ATP7A protein levels were not appreciably altered (Figure 2a).

**Figure 2.**
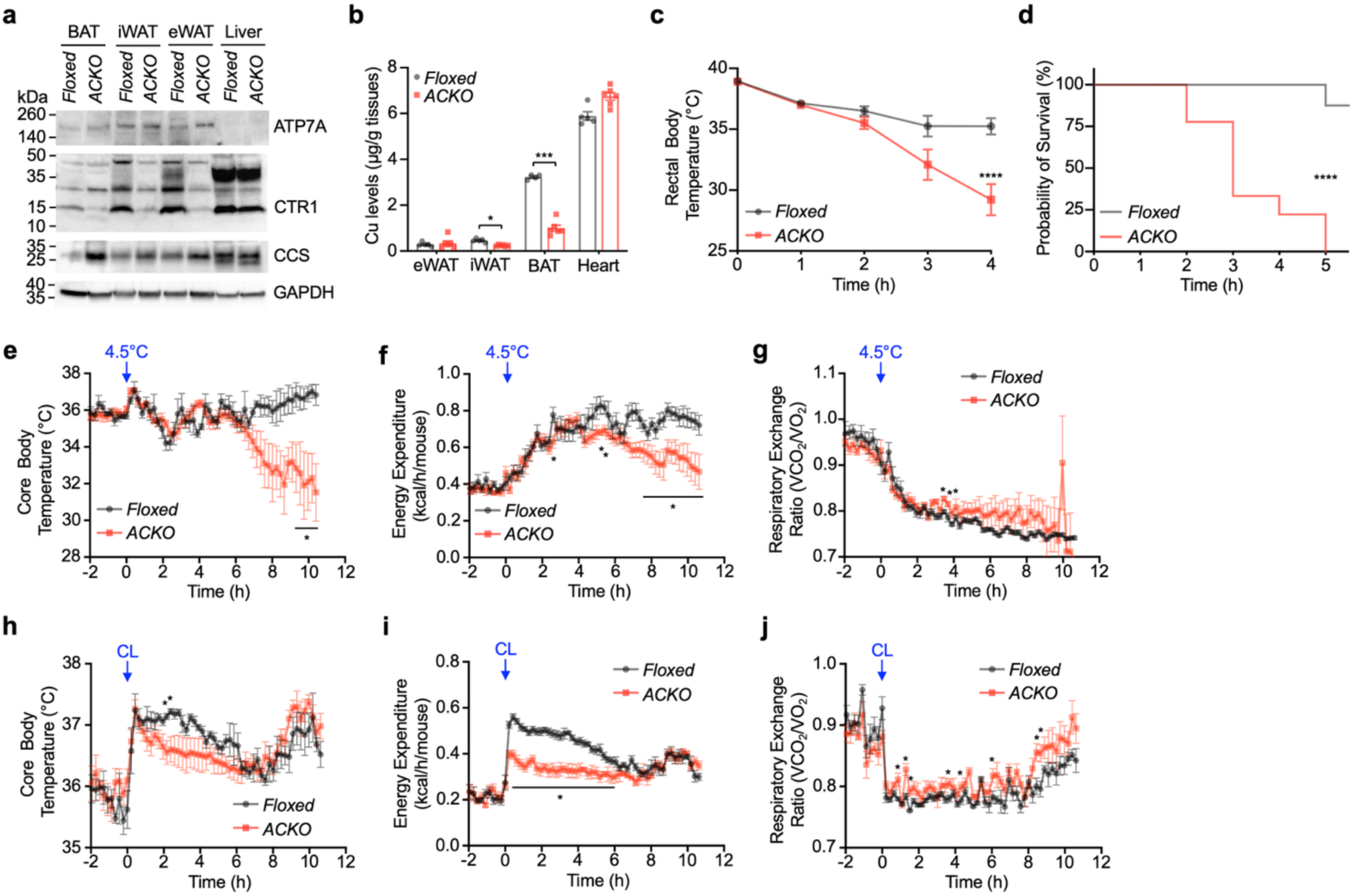
Adipocyte CTR1 is required for Cu homeostasis and thermogenic adaptation. (a) Immunoblot analysis of ATP7A, CTR1, and CCS in BAT, iWAT, eWAT, and liver from *Ctr1*-floxed (Floxed) and adipose-specific *Ctr1* knockout (ACKO; *Ctr1^fl/fl^*; *Adipoq*-Cre) mice. GAPDH served as a loading control. (b) ICP–MS quantification of Cu levels in tissues from Floxed (n = 5) and ACKO (n = 6) mice. (c) Rectal body temperature in Floxed (n = 6) and ACKO (n = 8) mice during acute cold exposure (CE, 4°C). Food, but not water, was removed at the onset of CE from RT (∼22°C). (d) Kaplan–Meier survival analysis during CE showing reduced survival in ACKO (n = 9) compared with Floxed (n = 8) mice (log-rank test, ****P < 0.0001). Mice were euthanized upon reaching the endpoint criterion (rectal temperature <28°C). (e–g) Indirect calorimetry measurements during gradual transition (∼80 min) from 22°C to 4.5°C. Food, but not water, was removed at the onset of cooling. Core body temperature (e), cold-induced energy expenditure (f), and respiratory exchange ratio (RER) (g) in Floxed (n = 6) and ACKO (n = 6) mice. (h–j) Core body temperature (h), CL-induced energy expenditure (i), and RER (j) before and after intraperitoneal injection of CL (1 mg kg⁻¹) in Floxed (n = 6) and ACKO (n = 6) mice. Data are presented as mean ± SEM. Statistical significance was determined by two-tailed Student’s t-test. *P < 0.05, **P < 0.01, ***P < 0.001, ****P < 0.0001 versus Floxed controls. Unless otherwise indicated, experiments were performed in ∼10-week-old male mice.

Under basal conditions at room temperature (RT), ACKO mice exhibited normal gross appearance and no significant differences in body weight, fat mass, lean mass, or adipose depot weights compared with floxed littermate controls (Figure S2c–g), indicating that adipocyte *Ctr1* deletion does not overtly disrupt adipose development or body composition under steady-state conditions.

In contrast, during acute cold exposure (4°C), ACKO mice exhibited a rapid and pronounced decline in rectal temperature (Figure 2c) and succumbed within 4–5 hours (rectal temperature <28°C, the humane endpoint), whereas floxed controls maintained body temperature and survived (Figure 2d). The severity of the ACKO phenotype exceeded that observed in dietary Cu-deficient WT mice (Figure 1c,d), supporting a critical cell-autonomous requirement for adipocyte CTR1 during thermogenic stress.

Consistent with impaired thermogenic activation^31, 32^, indirect calorimetry during a gradual temperature transition (∼80 min from 22°C to 4.5°C) revealed that ACKO mice exhibited reduced core body temperature, diminished cold-induced energy expenditure, and altered respiratory exchange ratio (RER) dynamics compared with floxed controls (Figure 2e–g). The more progressive temperature decline observed under metabolic cage conditions likely reflects partial thermal adaptation during gradual cooling to 4.5°C; nevertheless, ACKO mice remained markedly impaired in mounting an effective thermogenic response. Furthermore, adipocyte-specific deletion of *Ctr1* attenuated the stimulatory effects of CL on energy expenditure and core body temperature (Figure 2h–j). Together, these data demonstrate that adipocyte CTR1-mediated Cu import is required for full thermogenic activation during both acute cold exposure and β3-adrenergic stimulation.

### 2.3. CTR1-dependent Cu import supports mitochondrial remodeling and beige adipocyte recruitment

To define the molecular mechanisms underlying impaired thermogenic adaptation in *Ctr1*-deficient adipocytes, we performed quantitative proteomic analysis of BAT from control (Floxed) and adipose-specific *Ctr1* knockout (ACKO) mice following 6 h of cold exposure. Using a 1.5-fold cutoff and adjusted P < 0.05, 112 proteins were significantly differentially expressed in ACKO BAT (Figure 3a; Figure S3a; Supplementary Table 1). Among these, 83 downregulated proteins were enriched in metabolic pathways, including fatty acid metabolism, oxidative phosphorylation (OXPHOS), carbon metabolism, and thermogenesis (Figure 3a; Figure S3a). Protein–protein interaction and STRING analyses revealed coordinated suppression of mitochondrial and thermogenic modules in ACKO BAT (Figure S3b,c).

**Figure 3.**
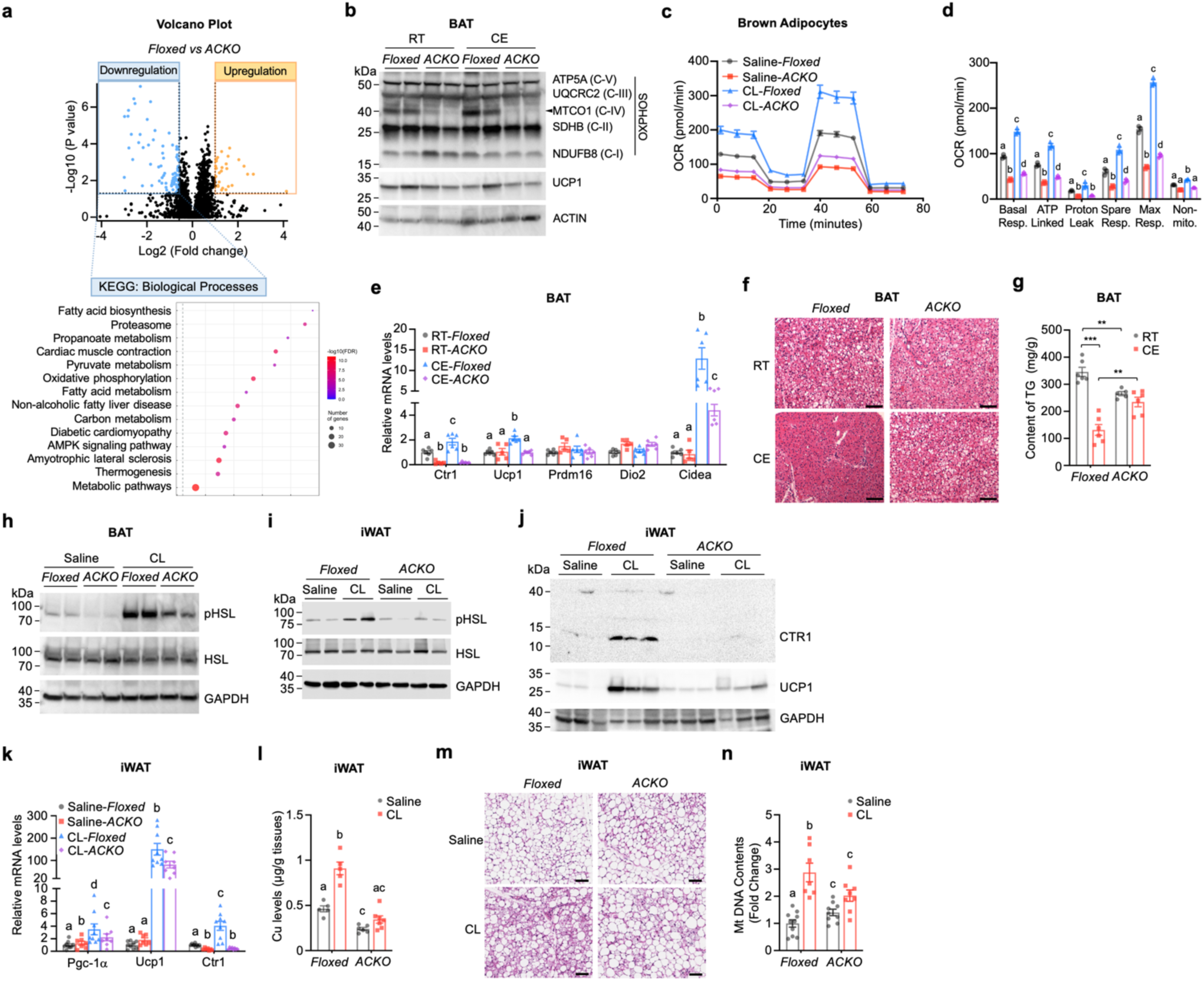
Adipocyte Cu deficiency impairs mitochondrial oxidative metabolism and lipid utilization in BAT and iWAT. (a) Volcano plot of differentially expressed proteins in BAT from Floxed and ACKO mice after 6 h CE (n = 3 per group). Downregulated proteins were subjected to KEGG pathway enrichment analysis (bottom). (b) Immunoblot analysis of OXPHOS complex subunits (ATP5A, UQCRC2, MTCO1, SDHB, NDUFB8), UCP1, and ACTIN in BAT from Floxed and ACKO mice housed at RT or exposed to cold (CE). (c) Oxygen consumption rate (OCR) traces of immortalized pre-brown adipocytes derived from Floxed and ACKO mice treated with vehicle or CL (10 μM). (d) Quantification of respiratory parameters derived from (c), including basal respiration, ATP-linked respiration, proton leak, maximal respiration, spare respiratory capacity, and non-mitochondrial respiration. (e) Relative mRNA expression of *Ctr1* and thermogenic genes (*Ucp1*, *Prdm16*, *Dio2*, *Cidea*) in BAT from Floxed and ACKO mice under RT or CE. (f) Representative H&E staining of BAT from Floxed and ACKO mice under RT or CE. Scale bar, 50 μm. (g) TG content in BAT from Floxed and ACKO mice under RT or CE. (h, i) Immunoblot analysis of phosphorylated HSL (Ser660) and total HSL in BAT (h) and iWAT (i) from Floxed and ACKO mice treated with saline or CL (1 mg kg⁻¹, 15 min). (j) Immunoblot analysis of CTR1 and UCP1 in iWAT from Floxed and ACKO mice treated with saline or CL. (k) Relative mRNA expression of *Ppargc1a*, *Ucp1,* and *Ctr1* in iWAT from Floxed and ACKO mice treated with saline or CL. (l) Cu content in iWAT from Floxed and ACKO mice treated with saline or CL. (m) Representative H&E staining of iWAT from Floxed and ACKO mice treated with saline or CL (1 mg kg⁻¹ day⁻¹, once daily for 10 consecutive days). Scale bar, 50 μm. (n) Relative mtDNA content in iWAT following 10 days of CL treatment. Data are presented as mean ± SEM. Statistical significance for panels (d, e, k, l, n) was determined by one-way ANOVA with Tukey’s post hoc test for each parameter analyzed independently. Groups not sharing a common letter are significantly different (P < 0.05).

Consistent with these findings, immunoblot analysis demonstrated a marked reduction in the Cu-dependent complex IV subunit MTCO1 in ACKO BAT under both RT and CE conditions (Figure 3b). Given that proper assembly and stability of MTCO1 within complex IV depend on mitochondrial Cu availability^33, 34^, the marked reduction in MTCO1 protein abundance indicates compromised Cu-dependent mitochondrial respiratory capacity. Functional assessment using immortalized pre-brown adipocytes derived from Floxed and ACKO mice showed that *Ctr1*-deficient cells exhibited significantly reduced oxygen consumption rates (OCR) at baseline and following β3-adrenergic stimulation with CL (Figure 3c,d), confirming impaired mitochondrial respiratory function. Together, these findings indicate that CTR1-dependent Cu import is required to sustain mitochondrial oxidative capacity during adrenergic activation.

We next examined thermogenic gene expression in vivo. Acute cold exposure robustly induced thermogenic transcripts in control BAT; however, this induction was attenuated in ACKO mice (Figure 3e). Notably, UCP1 protein abundance was comparable between genotypes (Figure 3b), indicating that the thermogenic impairment primarily reflects reduced mitochondrial oxidative capacity rather than diminished UCP1 protein levels.

We next assessed lipid utilization in BAT, since effective thermogenesis requires coordinated fatty acid mobilization through lipolysis to fuel mitochondrial respiration^35, 36^. Cold exposure reduced lipid droplet (LD) content in control BAT, whereas LD morphology was relatively preserved in ACKO BAT (Figure 3f). Triglyceride (TG) quantification confirmed impaired lipid clearance in ACKO BAT under CE conditions (Figure 3g). Moreover, CL–stimulated phosphorylation of hormone-sensitive lipase (HSL Ser660) was markedly attenuated in both BAT and iWAT of ACKO mice (Figure 3h,i), indicating defective β3-adrenergic–induced lipolytic signaling across adipose depots. These findings are consistent with prior observations that Cu inhibits PDE3B, thereby promoting cAMP-dependent PKA activation and lipolysis in adipocytes^18^ and suggest that impaired substrate mobilization may further constrain thermogenic output in ACKO mice.

Cu deficiency not only impairs BAT thermogenesis but also compromises beige adipocyte recruitment in white adipose tissue. Chronic β3-adrenergic stimulation robustly induced CTR1 and UCP1 protein expression in control iWAT, but this response was substantially blunted in ACKO mice (Figure 3j). Similarly, induction of thermogenic genes (*Ppargc1a* and *Ucp1*), as well as *Ctr1*, was significantly reduced in ACKO iWAT (Figure 3k). Importantly, CL-induced increases in Cu content in iWAT were markedly attenuated in ACKO mice (Figure 3l), indicating that adipocyte CTR1 is required for β3-adrenergic–stimulated Cu accumulation.

Histological analysis revealed reduced multilocular adipocyte formation in ACKO iWAT after chronic CL treatment (Figure 3m), accompanied by blunted mitochondrial DNA (mtDNA) expansion (Figure 3n), supporting a role for Cu in metabolic remodeling during adrenergic activation. To determine whether the observed thermogenic defects reflect adipocyte-intrinsic mitochondrial dysfunction, we examined isolated adipocyte cultures. Primary iWAT adipocytes derived from ACKO mice differentiated comparably to floxed controls (Figure S3d). However, upon β3-adrenergic stimulation with CL, ACKO adipocytes exhibited significantly reduced UCP1 induction compared to floxed controls (Figure S3e), demonstrating a cell-autonomous defect in the capacity of differentiated adipocytes to mount a thermogenic gene response.

Collectively, these findings demonstrate that adipocyte CTR1-mediated Cu import supports mitochondrial remodeling and thermogenic adaptation in both BAT and iWAT. In ACKO mice, impaired Cu-dependent mitochondrial function and defective lipolytic substrate mobilization together compromise the integrated adipose thermogenic response.

### 2.4. BAT-specific CTR1 deletion impairs Cu-dependent mitochondrial function and cold tolerance

To determine whether loss of *Ctr1* in BAT is sufficient to drive the cold-sensitive phenotype observed in ACKO mice, we generated BAT-specific *Ctr1* knockout mice (BCKO) using *Ucp1*-Cre–mediated deletion. CTR1 protein was efficiently depleted in BAT while remaining intact in iWAT (Figure S4a,b). Loss of CTR1 in BAT was accompanied by reduced abundance of the Cu-dependent complex IV subunit MTCO1 and CoxIV, whereas other examined OXPHOS components were less prominently affected, and mitochondrial protein levels in iWAT were largely unchanged. These findings are consistent with impaired Cu-dependent mitochondrial function selectively in BAT.

Under basal conditions, BCKO mice displayed normal body weight and adipose depot mass (Figure S4c–e), indicating preserved adipose tissue development. However, during acute cold exposure (4°C), BCKO mice rapidly developed hypothermia and reached the endpoint criterion (rectal temperature <28°C) compared with floxed controls (Figure S4f,g), demonstrating impaired thermogenic capacity. BAT histology following cold exposure revealed defective lipid remodeling (Figure S4h), consistent with reduced lipid utilization. LA–ICP–MS imaging further demonstrated diminished Cu signal intensity in BAT under both room temperature (RT) and cold exposure conditions, whereas Zn distribution was largely unchanged (Figure S4i), supporting a selective defect in Cu accumulation.

During gradual cooling from 22°C to 4.5°C in metabolic cages, BCKO mice exhibited reduced core body temperature accompanied by modest changes in energy expenditure and altered respiratory exchange ratio dynamics (Figure S4j–l), suggesting reduced thermogenic efficiency rather than global metabolic failure. Notably, following acute β3-AR stimulation with CL, BCKO mice mounted largely preserved acute thermogenic responses (Figure S4m–o). In contrast, pan-adipose *Ctr1* knockout mice showed markedly impaired responses (Figure 2h–j). BAT-specific *Ctr1* ablation largely recapitulated the cold-sensitive phenotype, establishing that Cu import within BAT is critical for sustained cold tolerance. However, ACKO mice exhibited more severe thermogenic impairment than BCKO mice, particularly in response to acute pharmacologic β3-adrenergic stimulation (Figure 2h-j vs Figure S4m-o). This differential sensitivity reveals a tissue-coordinated requirement for Cu across adipose depots. In ACKO mice, Cu deficiency in white adipose tissue impairs lipolytic signaling (reduced HSL phosphorylation, Figure 3h,i), limiting free fatty acid mobilization required to fuel BAT thermogenesis^37, 38^. In contrast, BCKO mice retain intact Cu-dependent lipolysis in iWAT, preserving substrate supply sufficient to support acute β3-adrenergic responses. However, prolonged cold exposure requires sustained mitochondrial oxidative capacity that depends critically on BAT Cu, explaining the preserved acute but impaired chronic cold tolerance in BCKO mice. Together, these findings suggest that CTR1-dependent Cu homeostasis supports multiple, functionally coupled components of the thermogenic cascade across adipose depots, including lipolytic activation and mitochondrial oxidative capacity.

### 2.5. Pharmacologic Cu delivery ameliorates thermogenic defects in adipocyte *Ctr1* knockout mice

To determine whether thermogenic impairment in ACKO mice is primarily driven by defective Cu delivery, we administered the Cu ionophore elesclomol (ES), which facilitates intracellular Cu transport independently of CTR1 and directs Cu to mitochondrial OXPHOS complexes^19, 39,40^. ACKO and *Ctr1*-floxed littermates received ES (10 mg kg⁻¹, once daily for three days) prior to acute cold exposure (4°C). As expected, ACKO mice rapidly developed hypothermia upon cold challenge. ES treatment attenuated the decline in core body temperature in ACKO mice, consistent with partial restoration of thermogenic capacity (Figure 4a).

**Figure 4.**
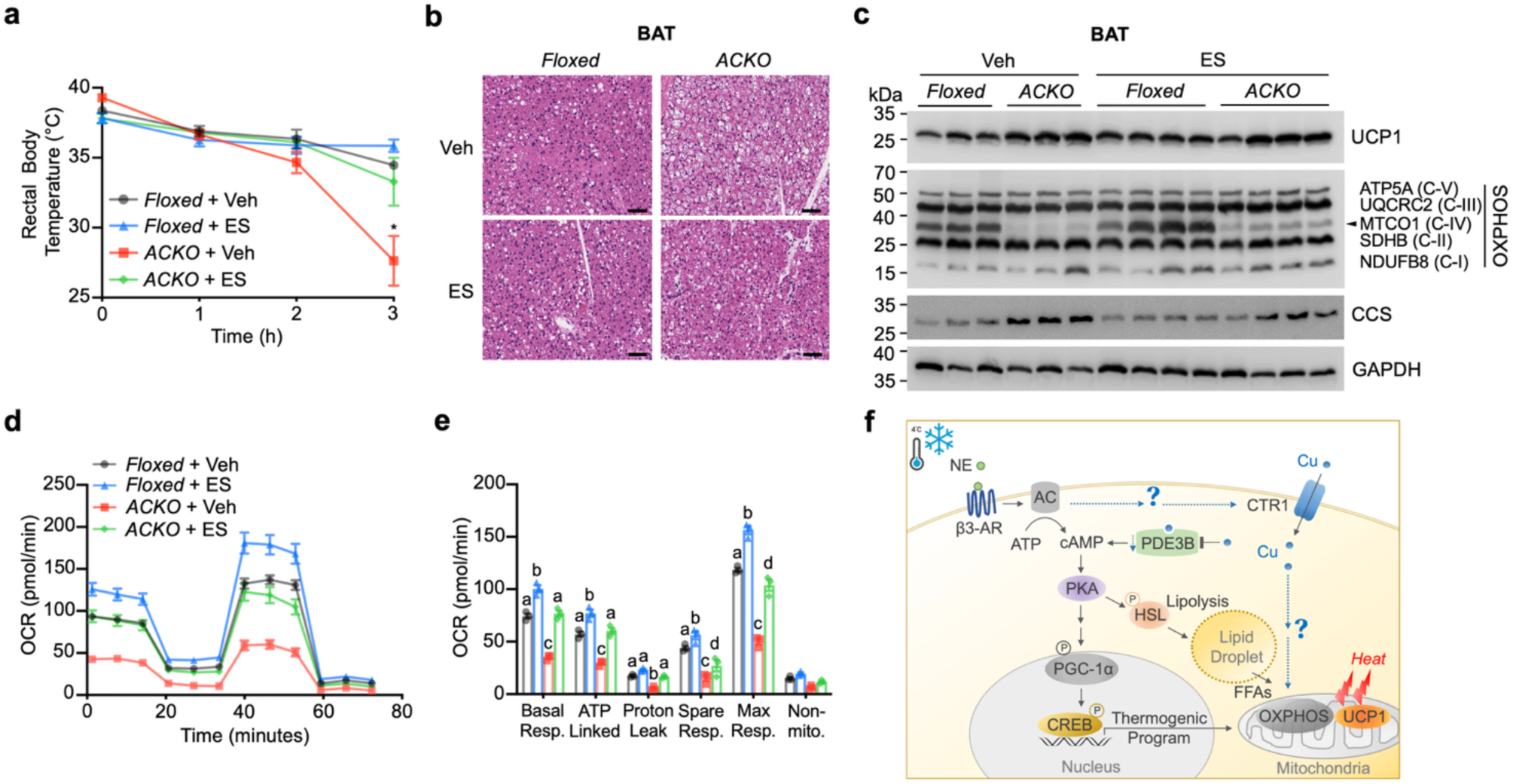
The Cu ionophore elesclomol rescues thermogenic defects in ACKO mice. (a) Rectal body temperature of *Ctr1*-floxed and ACKO mice treated with vehicle (Veh) or elesclomol (ES) during acute cold exposure (4°C). ES treatment improved cold tolerance (two-way ANOVA, treatment effect *P < 0.05; n = 3–4 per group). (b) Representative H&E staining of BAT from Ctr1-floxed and ACKO mice treated with Veh or ES under CE. Scale bar, 50 μm. (c) Immunoblot analysis of UCP1, OXPHOS complex subunits (ATP5A, UQCRC2, MTCO1, SDHB, NDUFB8), CCS, and GAPDH in BAT from *Ctr1*-floxed and ACKO mice treated with Veh or ES. (d, e) Mitochondrial respiration analysis in immortalized pre-brown adipocytes derived from *Ctr1*-floxed and ACKO mice treated with ES (10 nM) for 24 h. (d) Oxygen consumption rate (OCR) traces. (e) Quantification of basal respiration, ATP-linked respiration, proton leak, spare respiratory capacity, maximal respiration, and non-mitochondrial respiration. Data are presented as mean ± SEM. Statistical significance was determined by two-way ANOVA with Tukey’s post hoc test unless otherwise indicated. Groups not sharing a common letter are significantly different (P < 0.05, one-way ANOVA with Tukey’s post hoc test). (f) Working model illustrating the crosstalk between CTR1-mediated Cu import^56^ and adaptive thermogenesis^14, 57^. Upon cold exposure, norepinephrine (NE) activates β3-adrenergic receptor (β3-AR) signaling, increasing cAMP levels and activating protein kinase A (PKA). PKA promotes thermogenic gene expression (via PGC-1α and CREB) and phosphorylates hormone-sensitive lipase (HSL) to stimulate lipolysis. Released free fatty acids (FFAs) activate UCP1 and provide substrates for mitochondrial oxidative phosphorylation (OXPHOS). CTR1-dependent Cu import supports mitochondrial OXPHOS capacity and thermogenic output, whereas Cu delivery by elesclomol (ES) partially restores oxidative function in *Ctr1*-deficient adipocytes. The model highlights outstanding questions, including whether β3-AR signaling regulates CTR1 activity and whether mechanisms exist that prioritize Cu delivery to mitochondria during thermogenic activation.

Histological analysis revealed improved cold-induced BAT lipid remodeling in ES-treated ACKO mice compared with vehicle-treated ACKO controls (Figure 4b).

At the molecular level, ES treatment increased OXPHOS protein abundance in ACKO BAT, including partial restoration of the Cu-dependent complex IV subunit MTCO1, and reduced CCS accumulation, consistent with improved intracellular Cu availability (Figure 4c; Figure S5a). In *Ctr1*-deficient brown adipocytes, ES dose-dependently restored OXPHOS protein levels (Figure S5b) and enhanced mitochondrial respiration, as evidenced by increased basal and maximal oxygen consumption rates (Figure 4d, e). Notably, although ES substantially improved respiratory parameters in ACKO adipocytes, mitochondrial function was not fully normalized under the dosing regimen used in this study.

Collectively, these findings indicate that pharmacologic Cu delivery partially restores mitochondrial oxidative capacity and thermogenic function in the absence of CTR1, supporting a central role for intracellular Cu availability in adaptive thermogenesis.

## 3. DISCUSSION

Adaptive thermogenesis requires coordinated substrate mobilization, mitochondrial oxidative capacity, and transcriptional remodeling in adipose tissue. Although Cu is a well-established cofactor for mitochondrial enzymes such as cytochrome c oxidase, its dynamic regulation during thermogenic activation has remained unclear. Here, we identify CTR1-mediated Cu import as a regulated component of the β3-adrenergic thermogenic program that supports mitochondrial oxidative remodeling in thermogenic adipocytes.

Cold exposure and β3-adrenergic stimulation increased CTR1 abundance and Cu accumulation in thermogenic adipose tissues. Adipocyte-specific deletion of *Ctr1* reduced energy expenditure and caused cold intolerance, establishing a physiological requirement for Cu import in adaptive thermogenesis. Proteomic and functional analyses revealed suppression of oxidative phosphorylation and reduced mitochondrial respiratory capacity in *Ctr1*-deficient adipocytes, indicating compromised oxidative metabolism during thermogenic activation.

In addition to impaired oxidative metabolism, Cu deficiency was associated with attenuated cold- and β3-adrenergic–induced lipolytic activation in vivo, including reduced HSL phosphorylation and diminished lipid clearance. These observations are consistent with the proposed role of Cu as a direct metalloallosteric inhibitor of PDE3B in adipocytes^18^; Cu deficiency would be predicted to increase PDE3B-mediated cAMP degradation, thereby reducing PKA activity and HSL phosphorylation downstream of β3-AR signaling. Our findings are consistent with an extension of this Cu–PDE3B–cAMP regulatory axis to thermogenic adipocytes in a physiological context, suggesting that Cu availability contributes to the coordination of lipolytic activation during cold adaptation. In parallel, reduced mitochondrial oxidative capacity in *Ctr1*-deficient adipocytes may create a metabolic bottleneck that secondarily dampens lipolytic drive^41^, as impaired fatty acid oxidation limits the oxidative demand that normally sustains fatty acid mobilization from lipid droplets. Whether impaired cAMP–PKA signaling, reduced oxidative demand, or both underlie the attenuated lipolytic response in ACKO adipose tissue remains to be determined. Nonetheless, these findings suggest that Cu availability modulates both upstream cAMP signaling and downstream mitochondrial oxidative demand, thereby influencing multiple functionally coupled components of the thermogenic cascade across adipose depots.

BAT-specific *Ctr1* ablation largely recapitulated the cold-sensitive phenotype, suggesting that Cu import is required primarily within thermogenic adipocytes during physiological cold challenge. Notably, acute pharmacologic β3-adrenergic stimulation remained relatively preserved in BAT-specific knockout mice, whereas pan-adipose deletion impaired this response. Previous studies have shown that thermogenesis can be maintained despite impaired local BAT lipolysis^38^, provided that WAT supplies compensatory fuel through lipolysis and browning.

Together, these findings suggest that full thermogenic output, especially under acute demands for substrate mobilization, depends on coordinated, Cu-dependent functions across both brown and white adipose depots.

Partial restoration of thermogenic function by the Cu ionophore elesclomol further supports intracellular Cu availability as a limiting determinant of thermogenic competence. Together, these findings suggest that beyond its established role in mitochondrial enzyme assembly, Cu import appears to be dynamically upregulated during adrenergically driven metabolic remodeling.

Clinically, these findings may be relevant to Cu deficiency states. Hypothermia observed in Menkes disease and secondary Cu deficiency may reflect impaired thermogenic capacity ^15, 42, 43^. With the increasing prevalence of bariatric surgery and micronutrient malabsorption^44, 45^, understanding how trace metal homeostasis intersects with energy metabolism is increasingly important. Our results suggest that Cu availability represents a modifiable determinant of adipose thermogenic function.

Several mechanistic questions remain. These include how CTR1 abundance is regulated during adrenergic stimulation and whether its subcellular localization is dynamically remodeled; whether endosomal CTR1^46^ and associated Cu pools contribute to early thermogenic activation; and whether thermogenic activation selectively prioritizes mitochondrial Cu trafficking via Cu chaperones or a metalloadaptor^3^, and, if so, which Cu delivery machinery or mitochondrial import pathways, such as the SLC25A3 mitochondrial Cu transporter^47^, are engaged (Figure 4f). Addressing these questions will further clarify how trace metal homeostasis is coordinated with metabolic adaptation. Collectively, our study identifies intracellular Cu availability, and specifically CTR1-mediated Cu import, as a dynamically regulated determinant of adaptive thermogenesis that supports mitochondrial oxidative capacity and lipid mobilization to sustain effective cold adaptation.

### Limitations of the Study

Several limitations should be noted. Although our data establish a requirement for CTR1-dependent Cu import in thermogenic adipose tissue, the mechanisms linking β3-adrenergic signaling to the regulation of Cu import and intracellular distribution remain incompletely defined. Most mechanistic experiments were performed in male mice, and potential sex-dependent differences in Cu handling and thermogenic regulation warrant further study.

Although BAT-specific *Ctr1* deletion largely recapitulates the cold-sensitive phenotype, contributions from other adipose depots or adipose-resident cell populations cannot be fully excluded. The intracellular destination of cold- and β3-adrenergic–induced Cu accumulation, and the specific cuproenzymes that receive it, remain to be determined. Partial rescue of thermogenic function by the Cu ionophore elesclomol provides indirect evidence that mitochondria may represent a major site of increased Cu utilization; however, this pharmacologic approach may not fully recapitulate endogenous Cu trafficking dynamics, and direct quantification of mitochondrial Cu pools will require further investigation. Additional Cu-sensitive pathways beyond complex IV may also contribute to the observed phenotype. Finally, further studies are needed to determine how these mechanisms translate to thermogenesis in other animal species and in humans, and to clinical Cu deficiency states.

## 4. MATERIALS AND METHODS

### Animals and Experimental Interventions

Adipocyte-specific and brown adipocyte–specific *Ctr1* knockout mice were generated by crossing *Ctr1^flox/flox^*mice^48^ with *Adipoq*-Cre and *Ucp1*-Cre transgenic mice (The Jackson Laboratory, strains 028020 and 024670) to produce adipocyte-specific knockout (ACKO; *Ctr1^flox/flox^*; *Adipoq*-Cre) and BAT-specific knockout (BCKO; *Ctr1^flox/flox^*; *Ucp1*-Cre) mice, respectively. Age-matched *Ctr1^flox/flox^* littermates lacking Cre served as controls. Cre-mediated excision was confirmed by genomic PCR of adipose tissue, and loss of CTR1 protein was verified by immunoblotting.

All mice were maintained on a C57BL/6J background and housed at ∼22°C under a 12-h light/dark cycle with ad libitum access to food and water unless otherwise indicated. WT dietary Cu and WT cold/CL316,243 (CL) studies (Figure 1 and related supplementary data) were performed in mixed-sex cohorts with balanced male and female representation. Unless otherwise specified, subsequent experiments were conducted in ∼8–10-week-old male mice, as indicated in the figure legends.

### Dietary Cu Deficiency

To induce Cu deficiency, C57BL/6J dams were placed on a Cu-deficient purified diet (Teklad TD.80388) beginning at postnatal day 1 (P1). Control mice received the same base diet supplemented with 20 mg L⁻¹ CuSO₄ in drinking water^49^. Pups were weaned at P20 and maintained on the Cu-deficient diet for an additional 4 weeks after weaning (total ∼7 weeks) prior to experimentation.

### Cold Exposure and β3-Adrenergic Stimulation

For acute cold exposure, mice were single-housed and transferred from room temperature (∼22°C) to 4°C. Food, but not water, was removed at the onset of cold exposure. Rectal temperature was measured at the indicated time points using a thermal probe (Bioseb). Mice reaching a rectal temperature below 28°C were immediately euthanized to prevent hypothermia-induced organ damage. For chronic β3-adrenergic stimulation (browning), mice received CL (1 mg kg⁻¹, intraperitoneally) once daily for 10 consecutive days. For acute stimulation experiments, CL (1 mg kg⁻¹, i.p.) was administered as a single injection, and tissues were collected at the indicated time points (e.g., 15 min). For pharmacologic Cu delivery, mice were treated with the Cu ionophore elesclomol (ES; 10 mg kg⁻¹, subcutaneously) formulated in 0.5% Methocel™ (final 2% DMSO) once daily for 3 consecutive days prior to cold exposure. Vehicle-treated controls received the same formulation without ES. The elesclomol dose used (10 mg kg⁻¹) was selected based on prior studies demonstrating its efficacy in rescuing Cu deficiency phenotypes^19^. In vitro dose–response studies employed 1–10 nM elesclomol, based on prior cell culture studies^40, 50^. All animal procedures were approved by the Institutional Animal Care and Use Committees at the University of Maryland, Chonnam National University, and the National Institute of Diabetes and Digestive and Kidney Diseases.

### Metabolic Phenotyping and Body Composition

Body composition (fat mass and fat-free mass) was measured in non-anesthetized mice using a time-domain EchoMR-100H analyzer (Echo Medical Systems, Houston, TX). Indirect calorimetry was performed using the Oxymax/CLAMS system (Columbus Instruments, Columbus, OH). Mice were acclimated to metabolic chambers located within a thermoregulated enclosure for 2 days at 22°C. For cold exposure, the enclosure temperature was reduced from 22°C to 4.5°C beginning at 9:30 AM, with the ambient temperature gradually declining from 22°C to 4.5°C over 80 min. Food, but not water, was removed at the onset of cooling. Oxygen consumption (VO₂), carbon dioxide production (VCO₂), energy expenditure, respiratory exchange ratio (RER = VCO₂/VO₂), and body temperature were recorded. Core body temperature was continuously monitored using intraperitoneally implanted telemetry transmitters (Mini Mitter/Philips Respironics). The effect of acute administration of a β3-adrenergic agonist (CL, 1 mg kg⁻¹, intraperitoneally) on body temperature and energy expenditure was tested in mice adapted to 30°C for 24 h.

### Triglyceride Measurement

BAT triglyceride content was measured as previously described^51^. Tissue was homogenized in PBS, and lipids were extracted with chloroform:methanol (2:1). Extracted lipids were dried under nitrogen, dissolved in ethanol containing 25% Triton X-100, and triglycerides were quantified using a commercial kit (Asanpharm) and normalized to tissue weight.

### Immunoblotting

Tissues and cultured cells were lysed in ice-cold lysis buffer (PBS, pH 7.4, 1% Triton X-100, 0.1% SDS, and 1 mM EDTA) supplemented with protease inhibitors. Equal amounts of protein were resolved by SDS–PAGE and transferred to PVDF membranes. Primary antibodies included CTR1^52^, UCP1 (Abcam, ab23841), ATP7A (from Dr. S. Kaler’s laboratory^53^), CCS (Santa Cruz Biotechnology, sc-55561), OXPHOS cocktail (Abcam, ab110413), HSL (Abcam, ab45422), phospho-HSL (Ser660; Cell Signaling, #45804), CoxIV (Thermo Fisher Scientific, A21348), β-actin (Thermo Fisher Scientific, MA515739), and GAPDH (Thermo Fisher Scientific, MA515738). Blots were developed using SuperSignal™ substrates and quantified by densitometry using ImageJ.

### RNA and mtDNA Analysis

Total RNA was isolated using TRIzol and reverse-transcribed using ReverTra Ace (Toyobo). Quantitative PCR was performed using SYBR Green chemistry. Gene expression levels were normalized to *Rplp0* using the ΔΔCt method. For mitochondrial DNA content, genomic DNA was isolated, and mitochondrial *Cox2* was normalized to nuclear *Nrip1*.

### Histology

Adipose tissues were fixed in 4% paraformaldehyde, paraffin-embedded, sectioned (5 μm), and stained with hematoxylin and eosin (Histoserv, Inc). Images were acquired using brightfield microscopy.

### Primary and Immortalized Adipocyte Culture

Stromal vascular fractions (SVF) were isolated from BAT or iWAT of 2-week-old mice by collagenase digestion. Cells were cultured in DMEM containing 10% FBS and differentiated for 6–8 days using a standard beige adipogenic cocktail (insulin, dexamethasone, T3, indomethacin, IBMX, and rosiglitazone). For immortalization, pooled preadipocytes isolated from iWAT explants of *Ctr1^flox/flox^* mice were transduced with SV40 large T antigen^54^, and single-cell clones were established. Selected BAT and iWAT clones were transfected with a Cre-expressing plasmid to generate homozygous *Ctr1*-deleted (*Ctr1*⁻/⁻ or *Ctr1*-KO) lines. Immortalized *Ctr1*-KO preadipocytes were maintained in DMEM supplemented with 10% FBS.

### Mitochondrial Respiration Assay

Oxygen consumption rate (OCR) was measured using an XF96 Seahorse Analyzer (Agilent). Cells were seeded in XF96 plates and treated with CL (10 μM) or ES (1–10 nM, as indicated) for 24 h. Assay medium contained 1 mM pyruvate, 2 mM glutamine, and 10 mM glucose. Cells were equilibrated in a CO₂-free incubator for 1 h prior to measurement. OCR was recorded following sequential injections of oligomycin (1.5 μM), FCCP (1 μM), and rotenone/antimycin A (0.5 μM). Respiratory parameters were calculated according to the manufacturer’s guidelines and normalized to protein content.

### Inductively Coupled Plasma Mass Spectrometry (ICP-MS)

Tissues were digested in trace-metal grade nitric acid (HNO₃) at 90°C until fully dissolved, followed by the addition of hydrogen peroxide to ensure complete oxidation. Cu and Fe were quantified using a PerkinElmer DRC II ICP–MS equipped with a dynamic reaction cell to minimize polyatomic interferences. Calibration was performed using matrix-matched standards, and data were normalized to tissue wet weight.

### Laser Ablation Inductively Coupled Plasma Mass Spectrometry (LA-ICP-MS)

LA-ICP-MS was performed as described previously^55^. Tissues were embedded in OCT compound (Tissue-Tek), frozen in a dry ice/isopentane bath, and stored at −80 °C. Cryosections (20 μm) were ablated using an NWR213 laser with TV2 sample chamber (ESI) with the following parameters: 6 μm spot size, 2.3 J cm⁻² fluence, 15 μm s⁻¹ stage speed, 20 Hz repetition rate, 800 mL min⁻¹ helium carrier gas flow, and 6 μm pattern spacing. Ablated aerosol was analyzed for ^63^Cu and ^66^Zn using an iCAP-Qc ICP-MS (Thermo Fisher Scientific) with 0.4 s dwell time. Data were processed in Igor Pro using Iolite’s Trace Elements data reduction scheme in semi-quantitative mode with ^63^Cu or ^66^Zn as reference. Elemental concentrations were determined using matrix-matched standards.

### Quantitative Proteomics

BAT was harvested after 6 h of cold exposure (n = 3 per group). Proteins were extracted in 8 M urea buffer and digested using the filter-aided sample preparation (FASP) method. Each sample was reduced by incubation with Tris(2-carboxyethyl) phosphine (TCEP) at 37 °C for 30 min and alkylated with iodoacetic acid (IAA) at 25 °C for 1 h in the dark. After sequential washing with lysis buffer and 50 mM ammonium bicarbonate (ABC), the proteins were digested with trypsin (enzyme-to-protein ratio of 1:50, w/w) at 37 °C for 18 h. Then, trypsin was inactivated by acidification with 15 μL of formic acid (Honeywell, Charlotte, NC, USA). The digested peptides were desalted using C18 spin columns (Harvard Apparatus, Holliston, MA, USA), and the peptides were eluted with 80% acetonitrile containing 0.1% formic acid in water.

The samples were analyzed by nanoLC–MS/MS using a Q Exactive Orbitrap mass spectrometer coupled to an Ultimate 3000 system operating at 300 nL min⁻¹. The gradient of mobile phase was as follows: 4% solvent B for 14 min, 4–15% solvent B for 61 min, 15–28% solvent B for 50 min, 28–40% solvent B for 20 min, 40–96% solvent B for 2 min, holding at 96% solvent B for 13 min, 96–4% solvent B for 1 min, and 4% solvent B for 24 min. Ions were scanned in high resolution (70,000 for MS1 and 17,500 for MS2 at m/z 400), and the scan range was 400–2,000 m/z for both MS1 and MS2. Raw data were processed using Proteome Discoverer (v2.5) against the Mus musculus UniProt database. Proteins exhibiting a ≥1.5-fold change with an adjusted P < 0.05 (Benjamini–Hochberg correction) were considered significantly different.

### Statistical Analysis

Data are presented as mean ± SEM. Two-tailed unpaired Student’s t-tests were used for pairwise comparisons; Welch’s correction was applied when variances were unequal. One-way or two-way ANOVA followed by Tukey’s multiple comparisons test was used for multiple group comparisons as indicated. Kaplan–Meier survival curves were analyzed using the log-rank (Mantel–Cox) test. Analyses were performed using GraphPad Prism 10. P < 0.05 was considered statistically significant.

## ACKNOWLEDGEMENTS

We thank Daniel M. Lesperance, Sai Yuan, and Yinyan Ma for technical support and Phuc L. Nguyen for bioinformatic analysis of the proteomic data. We also thank the laboratory of Dr. Sarah L. J. Michel at the Metallotherapeutics Research Center, University of Maryland School of Pharmacy, for ICP–MS analysis. We are especially grateful to Dr. Stephen G. Kaler for providing the adipocytes from *mo-br* mice and the anti-ATP7A antibody and to Dr. Jaekwon Lee for helpful discussions. This work was supported by the National Research Foundation of Korea (RS-2021-NR058898 and RS-2023-00219517 to T.-I.J.), the National Institutes of Health Intramural Program (ZIC DK070002 to O.G.), the Maryland Agricultural Experiment Station (MAES grant #1763, FY21–22 to B.-E.K.), and the National Institutes of Health (GM79465 to C.J.C. and DK129599 to B.-E.K.).

## AUTHOR CONTRIBUTIONS

T.-I.J., and B.-E.K. designed the research; T.-I.J., Y.-S.L., T.K., J.K., P.P., P.B., X.Z., E.Y., N.L., and T.X. performed the research; T.-I.J., Y.-S.L., T.K., J.K., P.P., P.B., X.Z., E.Y., N.L., T.X., C.J.C., O.G., and B.-E.K. analyzed the data; T.-I.J., O.G., and B.-E.K. wrote the paper.

## DATA AVAILABILITY

The mass spectrometry proteomics data generated in this study have been deposited in the ProteomeXchange Consortium via the PRIDE partner repository under accession number PXD075401. All other data supporting the findings of this study are available from the corresponding authors upon reasonable request.

## APPENDIX A. SUPPLEMENTARY MATERIAL

Supplementary data are available.

