## Supplementary Material for "Copper Import via CTR1 Supports the β3-Adrenergic Thermogenic Program"

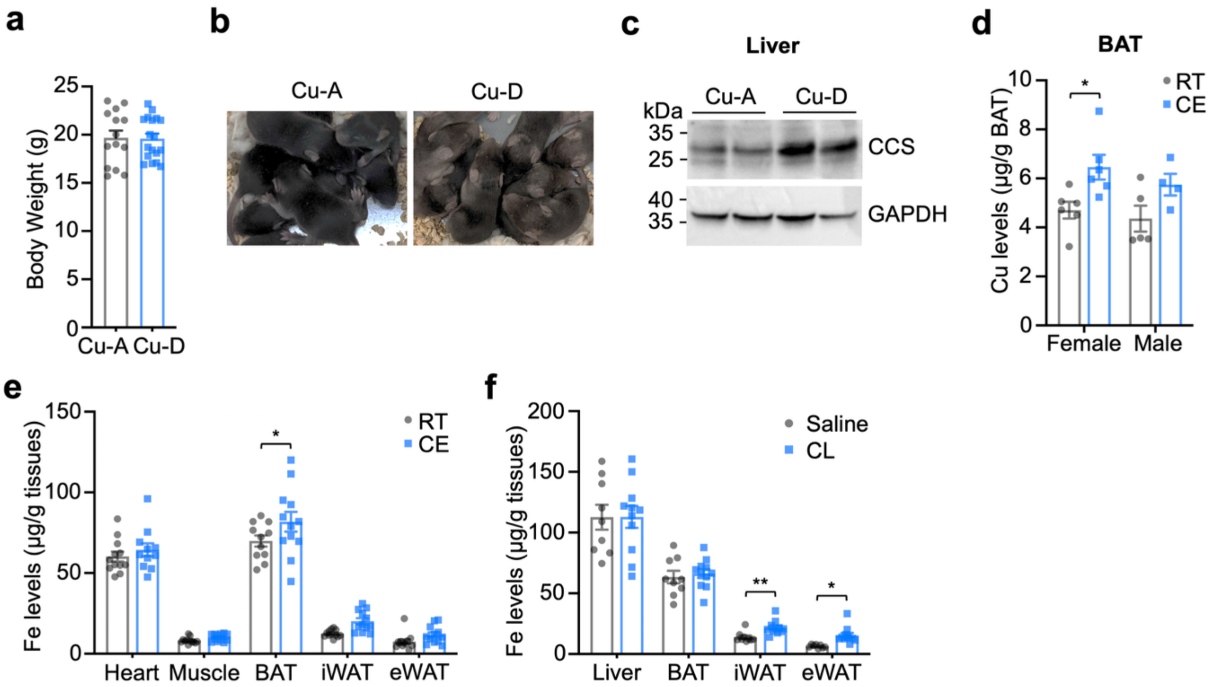


**Figure S1. Changes in tissue metal levels in cold-exposed (CE) or CL-treated WT mice.**
(a) Body weight of WT mice at the endpoint of dietary intervention (~7 weeks of age) maintained on Cu-adequate (Cu-A; n = 14; 7M, 7F) or Cu-deficient (Cu-D; n = 18; 9M, 9F) diets.
(b) Representative images of 10-day-old pups from Cu-D mothers showing a noticeably paler coat color compared with age-matched pups from Cu-A mothers.
(c) Immunoblot analysis of CCS protein levels in liver from Cu-A and Cu-D mice. GAPDH served as a loading control.
(d) ICP–MS quantification of Cu levels in BAT from male and female WT mice housed at RT or exposed to cold (CE, 4°C) for 12 h (n = 4–6 per sex per condition). Sex-stratified analysis revealed a significant cold-induced increase in BAT Cu in female mice (P = 0.0188) but a non-significant trend in males (P = 0.0846) (unpaired two-tailed Welch's t-test; RT vs CE within sex), suggesting possible sex-dependent differences in Cu handling during cold adaptation that warrant further investigation.
(e) ICP–MS analysis of iron (Fe) levels in heart, muscle, BAT, iWAT, and eWAT from WT mice housed at RT or CE (n = 9–12 per group).
(f) ICP–MS analysis of Fe levels in liver, BAT, iWAT, and eWAT from WT mice treated with saline or CL (n = 9–12 per group).
Unless otherwise indicated, mice were analyzed as mixed sex with balanced male and female representation across groups. Data are presented as mean ± SEM. Statistical significance was determined by two-tailed Student's t-test.

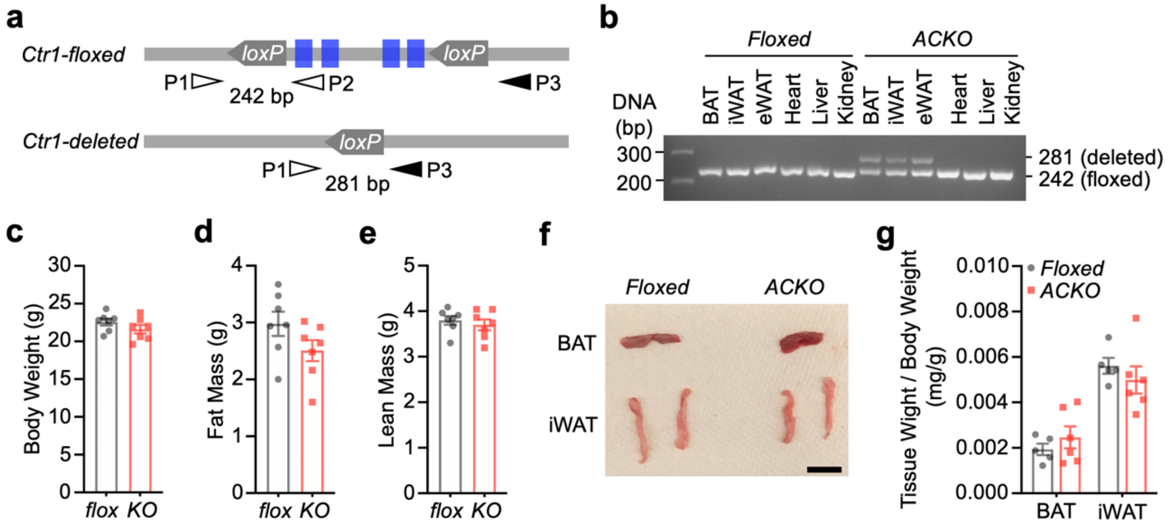


**Figure S2. Generation and body composition of adipose-specific *Ctr1* knockout (ACKO) mice.**

(a) Schematic of the strategy for adipose-specific deletion of *Ctr1*. *Ctr1*$^{fl/fl}$ mice were crossed with *Adipoq*-Cre transgenic mice to generate adipose-specific *Ctr1* knockout (ACKO; *Ctr1*$^{fl/fl}$; *Adipoq*-Cre) mice. The locations of PCR primers (P1, P2, and P3) are indicated by arrowheads. Blue boxes denote *Ctr1* exons. PCR products: 242 bp (*Ctr1*-floxed allele) and 281 bp (*Ctr1*-deleted allele).

(b) Representative genomic PCR analysis demonstrating tissue-specific Cre-mediated excision in 2-month-old Floxed and ACKO mice.

(c–e) MRI-based body composition analysis of 3-month-old Floxed and ACKO littermates showing body weight (c), fat mass (d), and lean mass (e) (n = 7 per group).

(f) Representative images of BAT and iWAT from 2-month-old Floxed and ACKO mice. Scale bar, 1 cm.

(g) BAT and iWAT mass normalized to body weight (mg tissue $g^{-1}$ body weight) in Floxed and ACKO mice (n = 5–6 per group).

Data are presented as mean ± SEM. Unless otherwise indicated, experiments were performed in ~8–10-week-old male mice.

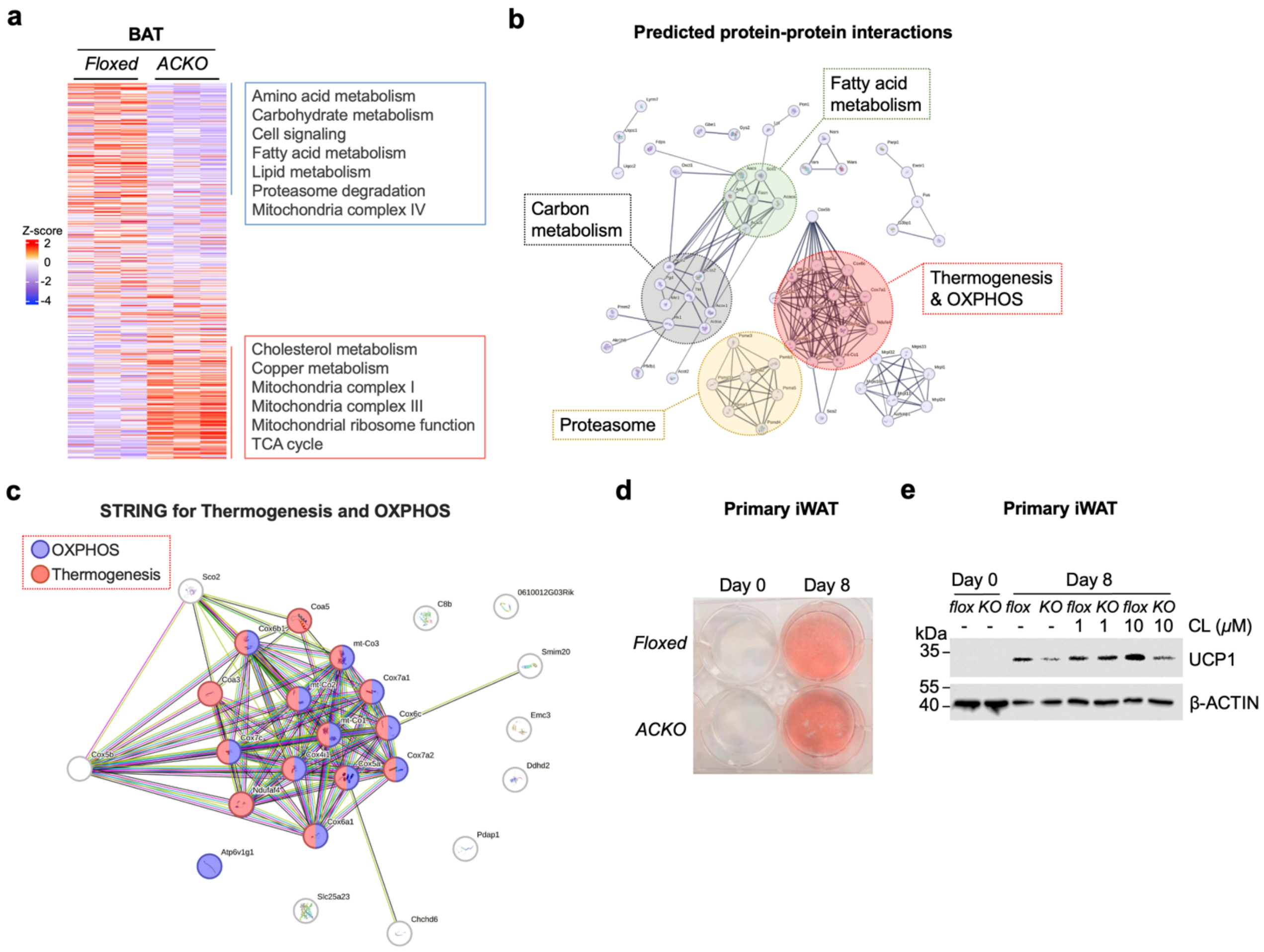


**Figure S3. Proteomic alterations and cell-autonomous defects in *Ctr1*-deficient adipocytes.**
(a) Heatmap of differentially expressed proteins in BAT from Floxed and ACKO mice after 6 h CE (n = 3 per group).
(b) Predicted protein–protein interaction network of downregulated proteins in ACKO BAT.
(c) STRING network of thermogenesis- and OXPHOS-related proteins reduced in ACKO BAT.
(d) Representative Oil Red O-stained images of differentiated primary iWAT adipocyte cultures (Day 0 and Day 8 after induction of differentiation) isolated from Floxed and ACKO mice.
(e) Immunoblot analysis of UCP1 in primary iWAT adipocytes derived from Floxed and ACKO mice treated with vehicle or CL (1 and 10 μM).

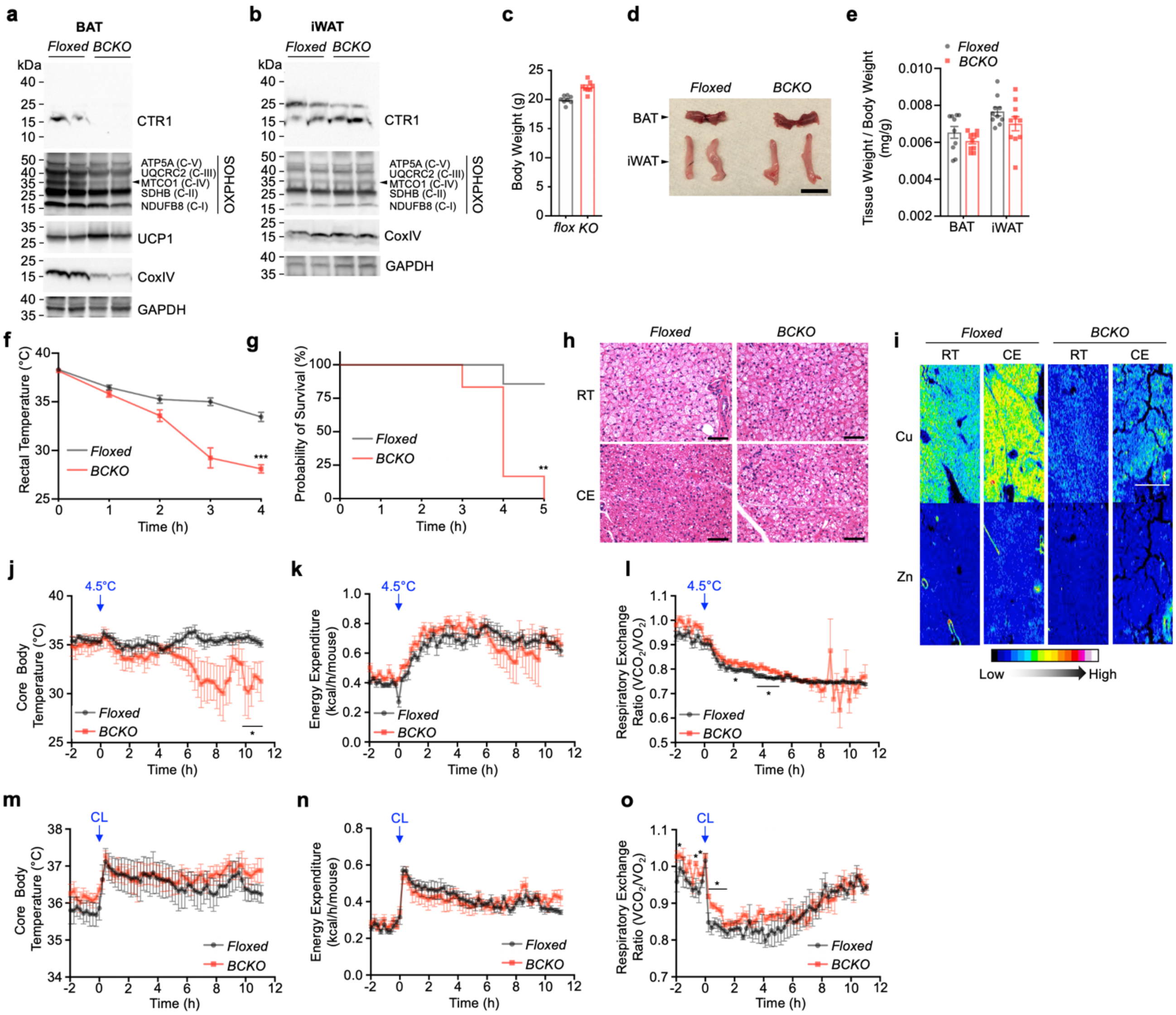

**Figure S4. BAT-specific *Ctr1* deletion causes cold intolerance and thermogenic defects.**
(a, b) Immunoblot analysis of CTR1, OXPHOS complex subunits (ATP5A, UQCRC2, MTCO1, SDHB, NDUFB8), UCP1 (BAT), CoxIV, and GAPDH in BAT (a) and iWAT (b) from *Ctr1*-floxed and BAT-specific *Ctr1* knockout (BCKO; Ucp1-Cre–mediated deletion) mice.
(c) Body weight of *Ctr1*-floxed and BCKO mice.
(d) Representative images of BAT and iWAT depots from *Ctr1*-floxed and BCKO mice.
(e) BAT and iWAT mass normalized to body weight in *Ctr1*-floxed and BCKO mice (n = 10 per group).
(f) Rectal temperature of *Ctr1*-floxed (n = 20) and BCKO (n = 17) mice during acute cold exposure (CE, 4°C). Mice were euthanized upon reaching the humane endpoint criterion.
(g) Kaplan–Meier survival curves of *Ctr1*-floxed (n = 6) and BCKO (n = 6) mice during CE (log-rank test, **P < 0.01). BCKO mice developed hypothermia more rapidly and reached the endpoint criterion (rectal temperature <28°C) sooner than controls.

(h) H&E staining of BAT from *Ctr1*-floxed and BCKO mice housed at RT or exposed to CE (3–4 h). Scale bar, 50 µm.
(i) LA–ICP–MS imaging of Cu and Zn distribution in BAT from *Ctr1*-floxed and BCKO mice under RT or CE (10 h). Scale bar, 500 µm.
(j–l) Indirect calorimetry analysis during transition from 22°C to 4.5°C in *Ctr1*-floxed (n = 6) and BCKO (n = 6) mice. (j) Core body temperature. (k) Total energy expenditure. (l) Respiratory exchange ratio (RER; $VCO_2/VO_2$).
(m–o) Metabolic responses following intraperitoneal injection of CL (1 mg $kg^{-1}$). (m) Core body temperature. (n) Total energy expenditure. (o) RER.
Data are presented as mean ± SEM. Statistical significance was determined by two-tailed Student's t-test (f, l, o) or two-way ANOVA with Tukey's post hoc test (j). *P < 0.05; **P < 0.01; ***P < 0.001; ****P < 0.0001. Unless otherwise indicated, experiments were performed in ~8–10-week-old male mice.

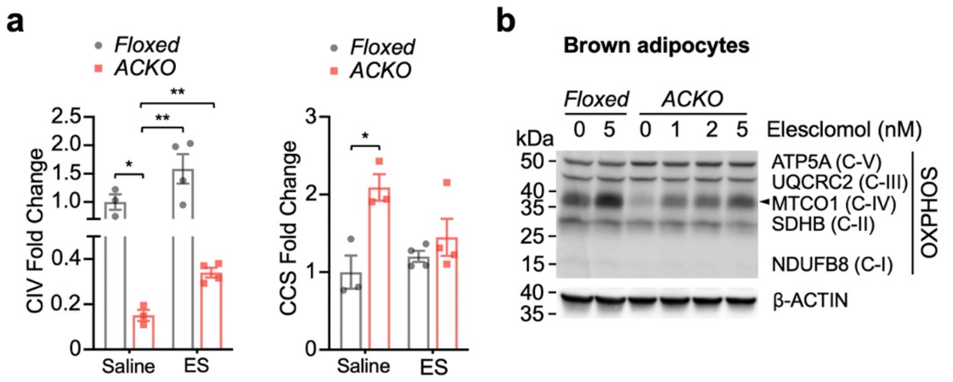


**Figure S5. Elesclomol restores mitochondrial complex IV and OXPHOS protein abundance in ACKO adipocytes.**

(a) Quantification of mitochondrial complex IV (CIV) and CCS protein levels in BAT from *Ctr1*-floxed and ACKO mice treated with vehicle or ES. Data are presented as fold change relative to floxed controls.

(b) Immunoblot analysis of OXPHOS complex subunits (ATP5A, UQCRC2, MTCO1, SDHB, NDUFB8) in brown adipocytes from *Ctr1*-floxed and ACKO mice treated with increasing concentrations of ES (nM) for 5 days. β-ACTIN served as a loading control.

Data are presented as mean ± SEM. Statistical significance was determined by two-way ANOVA with Tukey's post hoc test.
